# Highly specific intracellular ubiquitination of a small molecule

**DOI:** 10.1101/2024.01.26.577493

**Authors:** Weicheng Li, Enrique M. Garcia-Rivera, Dylan C. Mitchell, Joel M. Chick, Micah Maetani, John M. Knapp, Geoffrey M. Matthews, Ryosuke Shirasaki, Ricardo de Matos Simoes, Vasanthi Viswanathan, John L. Pulice, Matthew G. Rees, Jennifer A. Roth, Steven P. Gygi, Constantine S. Mitsiades, Cigall Kadoch, Stuart L. Schreiber, Jonathan M.L. Ostrem

## Abstract

Ubiquitin is a small, highly conserved protein that acts as a posttranslational modification in eukaryotes. Ubiquitination of proteins frequently serves as a degradation signal, marking them for disposal by the proteasome. Here, we report a novel small molecule from a diversity-oriented synthesis library, BRD1732, that is directly ubiquitinated in cells, resulting in dramatic accumulation of inactive ubiquitin monomers and polyubiquitin chains causing broad inhibition of the ubiquitin-proteasome system. Ubiquitination of BRD1732 and its associated cytotoxicity are stereospecific and dependent upon two homologous E3 ubiquitin ligases, RNF19A and RNF19B. Our finding opens the possibility for indirect ubiquitination of a target through a ubiquitinated bifunctional small molecule, and more broadly raises the potential for posttranslational modification *in trans*.

## Introduction

Until recently, the bulk of small-molecule drug development efforts focused on largely planar molecules having high sp^2^ hybridization (*1*), which might not optimally capitalize on the three-dimensional (3-D) complexity or stereospecificity of biological macromolecules. Diversity-oriented synthesis (DOS) produces small-molecule libraries whose members have diverse stereochemistry and 3-D skeletons. This has yielded novel chemical probes with diverse targets and functions (*2*, *3*). An innovative use of these libraries is to identify small molecules with entirely new modes of action within the cell through mechanistic studies of molecules with interesting phenotypes.

Mechanism of action (MOA) studies of small molecules have repeatedly led to important biological advances and entirely new avenues for drug development. MOA studies of the immunosuppressant natural products cyclosporin A and FK506 revealed for the first time that small molecules can function as molecular glues by inducing protein–protein interactions (*4*). Mechanistic studies revealed that the myeloma drug lenalidomide functions as a molecular glue degrader, co-opting the ubiquitin-proteasome system (UPS) and promoting ubiquitination and degradation of two zinc finger transcription factors, IKZF1 and IKZF3 by inducing their association with the E3 ubiquitin ligase cereblon (*5*, *6*). The discovery of molecular glues has led to extensive efforts to understand and harness the UPS for therapeutic applications. While many tools and drugs have been developed to modulate the UPS, none act at the level of ubiquitin itself.

### BRD1732 is a stereospecific cytotoxin dependent on the E3 ligases RNF19A and RNF19B

We identified the azetidine scaffold exemplified by BRD1732 from a DOS library as having stereospecific activity across multiple phenotypic screens. BRD1732 is broadly cytotoxic in cancer and immortalized cell lines, irrespective of lineage, with an IC_50_ of approximately 1 µM, and this effect is exclusive to the (2*S,3R,4R*) stereoisomer, meaning that altering one or more stereocenters on BRD1732 dramatically reduces activity (Fig. 1A and Fig. S1). Compared with BRD1732, both its enantiomer (BRD-E) and a diastereomer (BRD-D) are 10- to 50-fold less potent in viability assays across multiple cell lines. Our finding that activity depends on the 3-D orientation of the substituents on BRD1732 suggested to us that this molecule acts through direct interaction with one or more macromolecules.

**Fig. 1.**
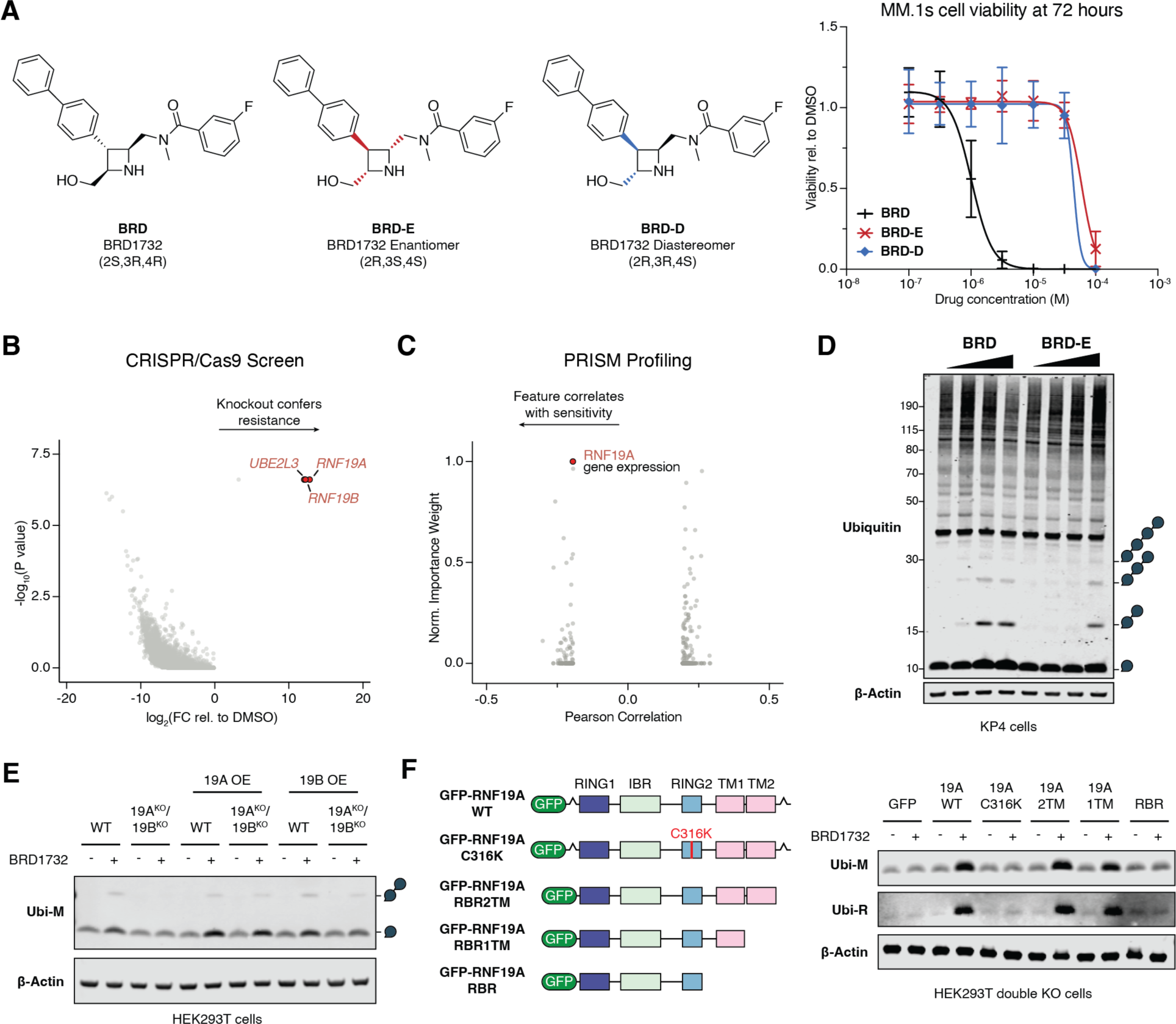
BRD1732 is a stereospecific cytotoxin dependent on the E3 ligases RNF19A and RNF19B. (**A**) Structures of BRD1732 (BRD), its enantiomer (BRD-E), and a diastereomer (BRD-D). Cell viability after 72-hour drug treatment of BRD, BRD-E, and BRD-D, in MM.1S cells. Fit was generated by 4-variable non-linear regression. (**B**) Genome-wide CRISPR/Cas9 knockout screen in MM.1S^Cas9^ multiple myeloma cells treated with 5 µM BRD1732 or DMSO. Top hit genes enriched in BRD1732-treated cells are highlighted in red. (**C**) PRISM screen of pooled, barcoded cancer cell lines treated with 5 µM BRD1732 showing sensitivity feature Pearson correlation with AUC. Importance weight calculated using enet. (**D**) Immunoblots of lysate from KP4 cells treated with BRD1732 and BRD-E for 6 hours at 0, 1, 10, and 100 µM. Ubiquitin chain symbols indicate the expected molecular weight for unanchored ubiquitin chains of various lengths. (**E**) Immunoblot of HEK293T WT and RNF19A/B double knockout cells transfected with GFP, RNF19A or RNF19B then treated for 6 hours with 5 µM BRD1732 or DMSO. (**F**) Schematic diagram of RNF19A mutants and truncation constructs. Immunoblots of HEK293T double KO cells transfected with the indicated expression constructs then treated for 6 hours with 5 µM BRD1732 or DMSO.

To identify the molecular machinery required for the cytotoxicity of BRD1732, we performed a genome-wide CRISPR-Cas9 resistance screen (Fig. 1B and Data S1). sgRNAs targeting *RNF19A*, *RNF19B* and *UBE2L3* were significantly enriched following BRD1732 treatment, suggesting that loss of these genes confers resistance. RNF19A and RNF19B are homologous RING-in-between-RING (RBR) E3 ubiquitin ligases, and UBE2L3 is an E2 ubiquitin ligase responsible for activating the RNF19 proteins (*7*). Using the PRISM platform (*8*) to profile the dependency features of BRD1732 across ∼580 cancer cell lines, we identified RNF19A expression as a feature strongly associated with sensitivity to BRD1732 (Fig. 1C). Taken together, these bidirectional findings strongly support a central role of RNF19A/B in the cytotoxicity of BRD1732.

While further interrogating the effects of BRD1732 on the UPS, we made the unexpected discovery that treatment of cells with BRD1732 causes accumulation of monoubiquitin and short polyubiquitin chains. Importantly, this effect is concentration-dependent and stereospecific, with EC_50_ values for BRD1732 and BRD-E approximating their corresponding IC_50_ values in viability assays (Fig. 1D). To understand whether this accumulation of ubiquitin is dependent on RNF19 proteins, we performed CRISPR-Cas9 knockout of both RNF19A and RNF19B in HEK293T cells.

RNF19A/B double knockout (DKO) cells showed no accumulation of mono- or di-ubiquitin upon treatment with BRD1732. Re-expression of RNF19A or RNF19B in DKO HEK293T cells is sufficient to recover production of these ubiquitin species (Fig. 1E). RBR E3 ligases require transfer of ubiquitin from the E2 ligase to a catalytic cysteine in the E3 RING2 domain prior to transfer to a substrate. Importantly, BRD1732-induced accumulation of ubiquitin requires the RBR core of RNF19A (RING1-IBR-RING2), including the catalytic cysteine C316, and at least one of its two transmembrane domains, but it does not require the N-terminal extension or the C-terminal regulatory domain (Fig. 1F).

### BRD1732 is directly ubiquitinated in cells

In addition to accumulation of low molecular weight polyubiquitin chains, we noticed that BRD1732 induces a very small but reproducible gel shift for monoubiquitin, an effect that is even more prominent on 12% Bis-Tris polyacrylamide gel (Fig. 1D and Fig. S2). We reasoned that this could indicate a change in posttranslational modification of ubiquitin or conjugation of ubiquitin to a small molecule or metabolite. BRD1732-induced ubiquitin accumulation is more pronounced in cells overexpressing RNF19 (Fig. 1E and 1F), so we purified endogenous ubiquitin from RNF19A-overexpressing Expi293F cells either untreated or treated with 10 µM BRD1732 for 6 hours. We then purified these ubiquitin samples by cation exchange followed by size exclusion chromatography and analyzed them by liquid chromatography–mass spectrometry (LC-MS) (Fig. 2A). The mass of endogenous ubiquitin from untreated cells matches the calculated mass for unmodified ubiquitin (8565 Da). However, the mass of endogenous ubiquitin from BRD1732-treated cells is 387 Da greater, which exactly matches the expected mass of a covalent adduct between ubiquitin and BRD1732 with loss of water (8952 Da).

**Fig. 2.**
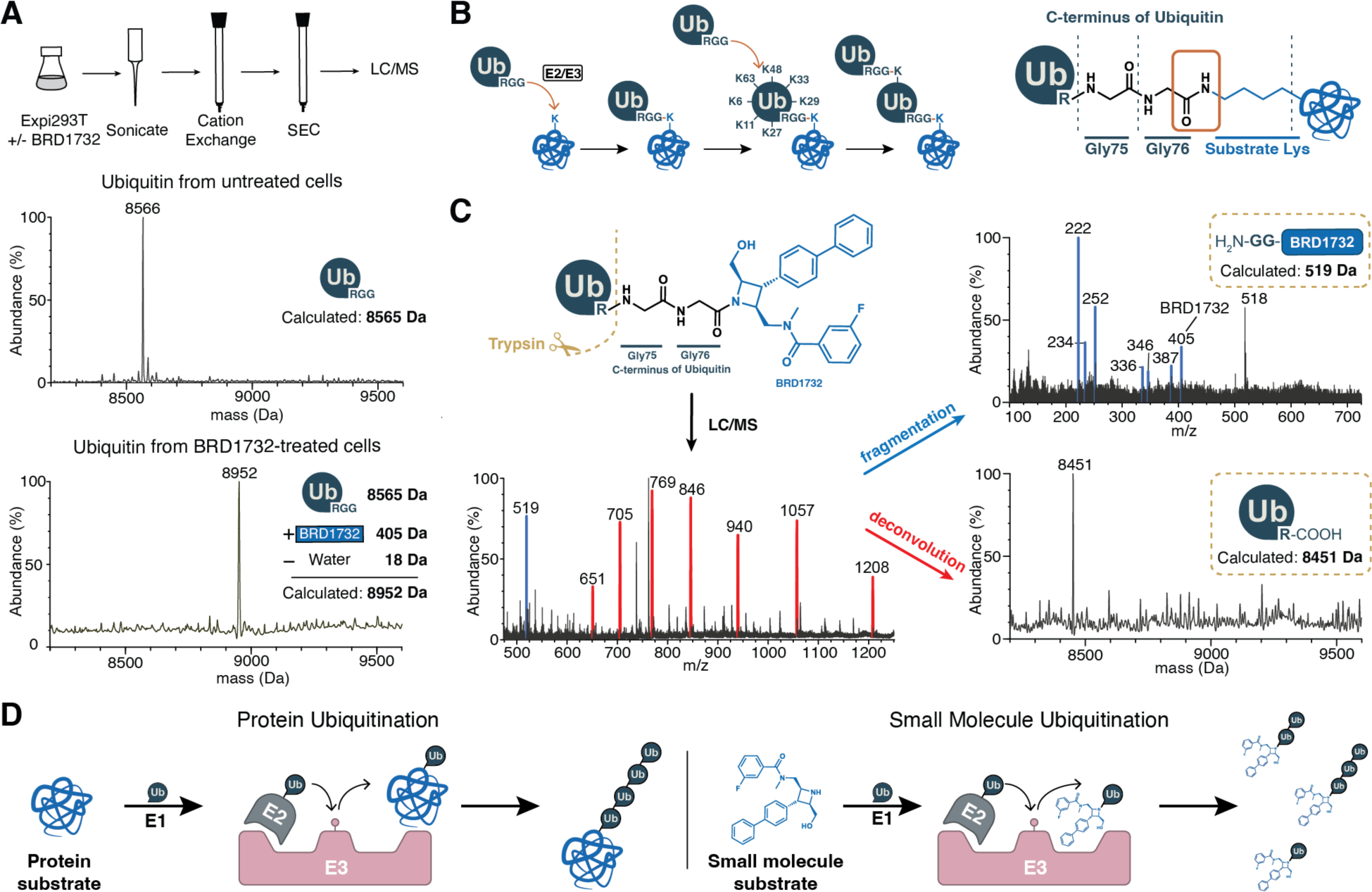
BRD1732 is directly ubiquitinated in cells. (**A**) Schematic of experiment workflows of purifying and characterizing the ubiquitin proteins from untreated or BRD1732-treated Expi293T cells (top). Deconvoluted mass spectra of intact LC-MS of purified ubiquitin from untreated and BRD1732-treated cells (bottom). (**B**) Schematic representations conjugation of ubiquitin C-terminal glycine and ε-amino group of a lysine on the target substrate. (**C**) Schematic representation of trypsin digestion of ubiquitin-BRD1732 conjugate with intact protein LC-MS spectrum of trypsin-digested ubiquitin from BRD1732-treated (left). LC-MS-MS spectrum from fragmentation of the 519 ion and spectrum from deconvolution of the envelope of peaks highlighted in red are shown (right). (**D**) Schematic representation of protein substrate ubiquitination by ubiquitin ligases along with proposed mechanism of small molecule ubiquitination.

Ubiquitin is transferred to substrate lysine residues by formation of an isopeptide bond between the C-terminus of ubiquitin and the ε-amine of lysine resulting in loss of a water molecule (Fig. 2B) (*9*). Given that BRD1732 contains a free aliphatic amine like lysine, we suspected that BRD1732 was mimicking a substrate protein and undergoing direct ubiquitination (Fig. 2C). To characterize further the ubiquitin conjugate formed in cells, we performed trypsin digestion under nondenaturing conditions. Digestion of unmodified ubiquitin under these conditions results in a single cleavage after Arg74 to yield ubiquitin(1-74), as previously reported (*10*), with a mass of 8452 Da by LC-MS (Fig. S3). Similar trypsin digestion of the BRD1732-induced ubiquitin conjugate yielded ubiquitin(1-74) as well as a new fragment ion at m/z = 519, which corresponds to the predicted mass of Gly-Gly-BRD1732 ([M+H]^+^ = 519 Da). Further fragmentation of this 519 ion yielded BRD1732 ([M+H]^+^ = 405) and its expected fragmentation ions (Fig. 2C). Taken together, these results indicate that BRD1732 is directly ubiquitinated by attachment to the ubiquitin C-terminus in cells in a manner that is dependent upon the E3 ligases RNF19A and RNF19B and similar to the ubiquitination of protein substrates (Fig. 2D).

### BRD1732 disrupts the ubiquitin-proteasome system at multiple pathway nodes

Following purification of endogenous ubiquitin from BRD1732-treated cells, we exclusively detect the ubiquitin-BRD1732 conjugate by LC-MS. Therefore, we wondered whether BRD1732 treatment might deplete free ubiquitin in the cell. Indeed, treatment of wild type HEK293T or KP4 cells with BRD1732 caused concentration-dependent loss of histone H2A lysine-119 ubiquitination (Ub-H2A-K119), suggesting depletion of nuclear ubiquitin (Fig. 3A). Because loss of H2A-K119 ubiquitination can also indicate more general ubiquitin-related stress, we directly quantified free monoubiquitin by flow cytometry staining with HA-tUI (Fig. 3B) (*11*). HA-tUI is a tagged synthetic protein designed to bind free ubiquitin with high affinity (dissociation constant 66 pM) and high specificity. Using this method, HEK293T cells treated with BRD1732 showed a significant decrease in free ubiquitin compared with DMSO-treated cells. However, we note that this effect does not show a clear dose-response. Taken together these effects did not adequately explain the toxicity of BRD1732.

**Fig. 3.**
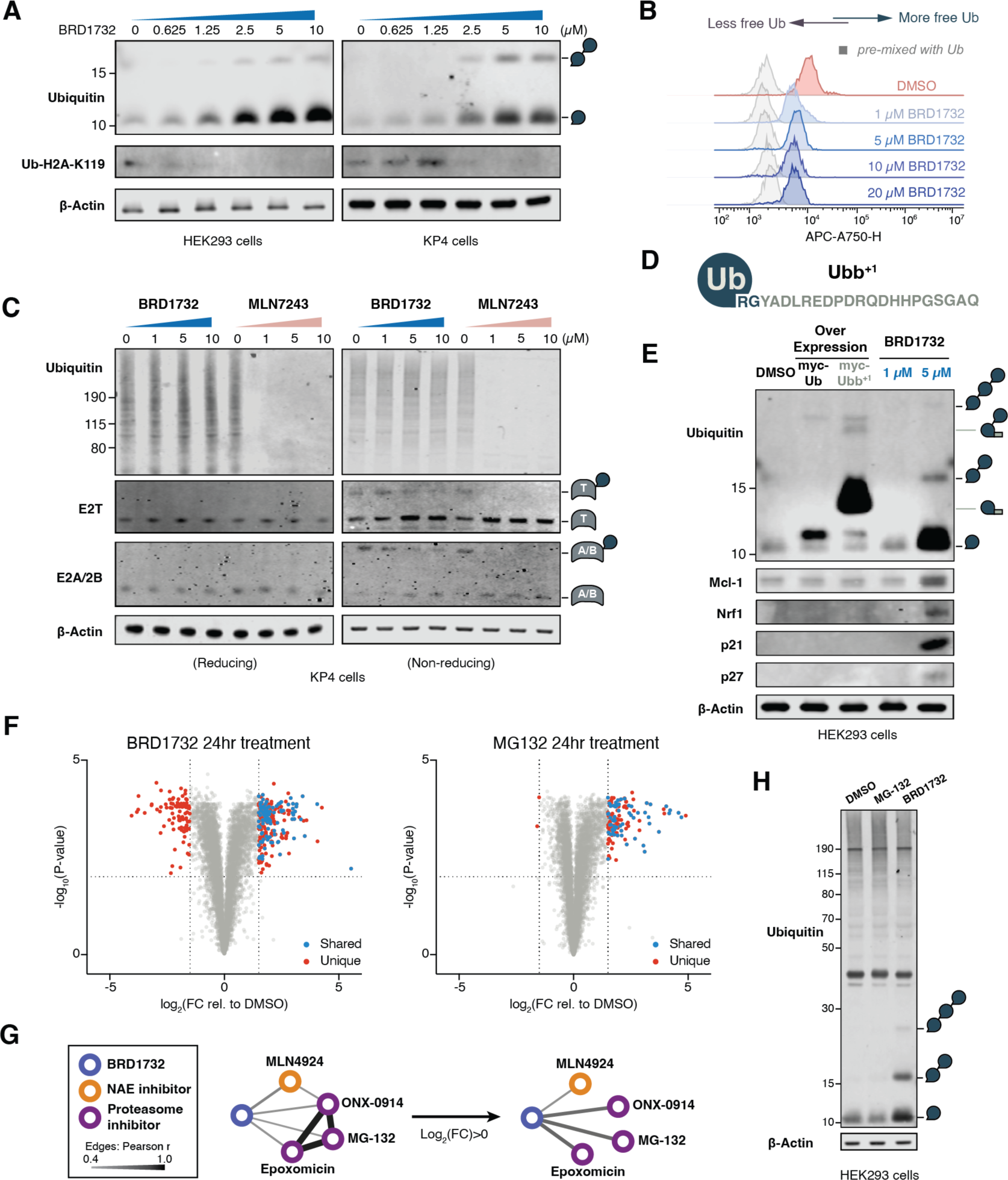
BRD1732 disrupts the ubiquitin-proteasome system at multiple pathway nodes. (**A**) Immunoblots following treatment of HEK293T or KP4 cells with BRD1732 at indicated concentrations for 24 hours. (**B**) Flow cytometry of HEK293T cells treated with BRD1732 at indicated concentrations for 6 hours, stained with free-ubiquitin binder HA-tUI followed by Alexa Fluor 750-conjugated anti-HA antibody. (**C**) Immunoblot following treatment of KP4 cells with indicated concentrations of BRD1732 or MLN7243 for 6 hours. (**D**) Schematic representation of UBB^+1^ with 20-amino acid C-terminal extension. (**E**) HEK293T cells were treated with BRD1732 for 24 hours or were transfected with myc-tagged ubiquitin or myc-tagged UBB+1. Accumulation of polyubiquitin and stabilization of short half-life proteins were analyzed by immunoblotting. (**F**) Yamato synovial sarcoma cells were treated with 5 µM BRD1732 or 5 µM MG-132 for 24 hours, then analyzed by TMT quantitative proteomics. Plot shows only proteins with no significant change in corresponding gene expression by RNA-Seq. Proteins significantly increased or decreased by both BRD1732 and MG132 are highlighted in blue. Proteins that change significantly in response to only one of these treatments are highlighted in red. (**G**) Compound similarity by quantitative proteomics in HCT116 cells. Network edge thickness corresponds to Pearson *r*. All nodes and network edges higher than 0.4 (Pearson *r*) are shown. (**H**) Immunoblots of 24-hour drug treatment of DMSO, MG132, and BRD1732 in HEK293T cells.

The adenosine monophosphate (AMP) mimetic ubiquitin activating enzyme (UAE) inhibitor MLN7243 conjugates to the C-terminus of ubiquitin within the UAE active site (*12*). The resulting ubiquitin-MLN7243 conjugate potently inhibits the UAE itself. Given the broad requirement of UAE for ubiquitination within the cell, MLN7243 treatment results in depletion of all conjugated and unconjugated ubiquitin species with the exception of free monoubiquitin (Fig. 3C, Fig. S4). As expected, charging of E2 ligases by UAE is also prevented by MLN7243. In contrast, BRD1732 only partially blocks UAE activity as indicated by ubiquitin charging of the E2 ligases E2T and E2A/2B, and BRD1732 has minimal effect on high molecular weight polyubiquitin conjugates at concentrations of up to 10 μM (Fig. 3C). These results are consistent with BRD1732 having a mechanism that is distinct from the UAE inhibitor MLN7243.

The terminal diglycine of ubiquitin is required for activation by ubiquitin ligases, conjugation to substrate proteins, and removal by deubiquitinases (*10*, *13*, *14*). In neurodegenerative diseases, molecular misreading results in a frameshift mutation in the ubiquitin B (*UBB*) gene resulting in production of an aberrant form of ubiquitin with Gly-76 replaced by a 20 amino acid extension (Fig. 3D) (*15*). Unanchored polyubiquitin chains terminating in UBB^+1^ have been reported to inhibit proteasomal degradation in cells, and we wondered whether ubiquitin-BRD1732 conjugates could behave similarly (*16*). While treatment with BRD1732 induced accumulation of multiple proteins known to undergo rapid proteasomal degradation, including MCL1, NRF1, p21 and p27, these proteins were not stabilized by overexpression of myc-tagged UBB^+1^ (Fig. 3E).

Given the striking effects on these proteins, we wondered whether BRD1732 might be more broadly affecting protein homeostasis. To assess this, we performed quantitative proteomics comparing BRD1732 with a proteasome inhibitor, MG-132, in Yamato synovial sarcoma cells (Fig. 3F and Data S2-S3). Focusing on proteins with no significant gene expression changes upon treatment by RNA-Seq (log_2_FC < 1, *p*_adj_ < 0.01), we identified 138 proteins that were stabilized by the proteasome inhibitor MG-132 and only 2 proteins that were destabilized. In contrast, BRD1732 induced dramatic bidirectional changes in protein stability, with 225 proteins significantly increased and 94 proteins significantly decreased. Strikingly, 63 proteins were stabilized by both BRD1732 and MG-132.

In HCT116 colorectal cancer cells, BRD1732-induced protein level changes correlate most strongly with proteasome inhibitors MG-132, epoxomicin, and ONX-0941 (Pearson coefficient *r* = 0.4-0.43), and the NEDD8-activating enzyme (NAE) inhibitor MLN4924 (*r* = 0.48, Fig. 3G and Data S4) when compared with 877 drugs and tool compounds reported in the same cell line as part of DeepCoverMOA (*17*). Restricting this analysis to proteins that increase upon treatment with BRD1732, we find that correlation with proteasome inhibitors strengthens (*r* = 0.58-0.6). However, unlike BRD1732, MG-132 does not induce accumulation of low molecular weight polyubiquitin chains but rather leads to accumulation of high molecular weight polyubiquitinated substrate proteins (Fig. 3H). These data suggest that BRD1732 may act in part through inhibition of the proteasome, but with a mechanism that is distinct from known proteasome inhibitors.

### BRD1732 disrupts ubiquitin-dependent proteasomal degradation

To understand the early effects of BRD1732 on protein homeostasis, we assessed proteome changes following 1-, 2-, and 4-hour treatments in HCT116 cells. Upon treatment with BRD1732, we observed a rapid and significant reduction in the levels of two highly homologous zinc finger proteins, ZFAND5 and ZFAND6 (Fig. 4A and Data S4). Co-treatment with MG-132 prevents loss of ZFAND5/6 suggesting that BRD1732 induces their degradation by the proteasome (Fig. 4B). Surprisingly, however, this effect is not prevented by the UAE inhibitor MLN7243, indicating that ZFAND5/6 undergo ubiquitin-independent proteasomal degradation. Likewise, ZFAND5/6 are efficiently degraded in cells lacking RNF19A and RNF19B (Fig. 4C).

**Fig. 4.**
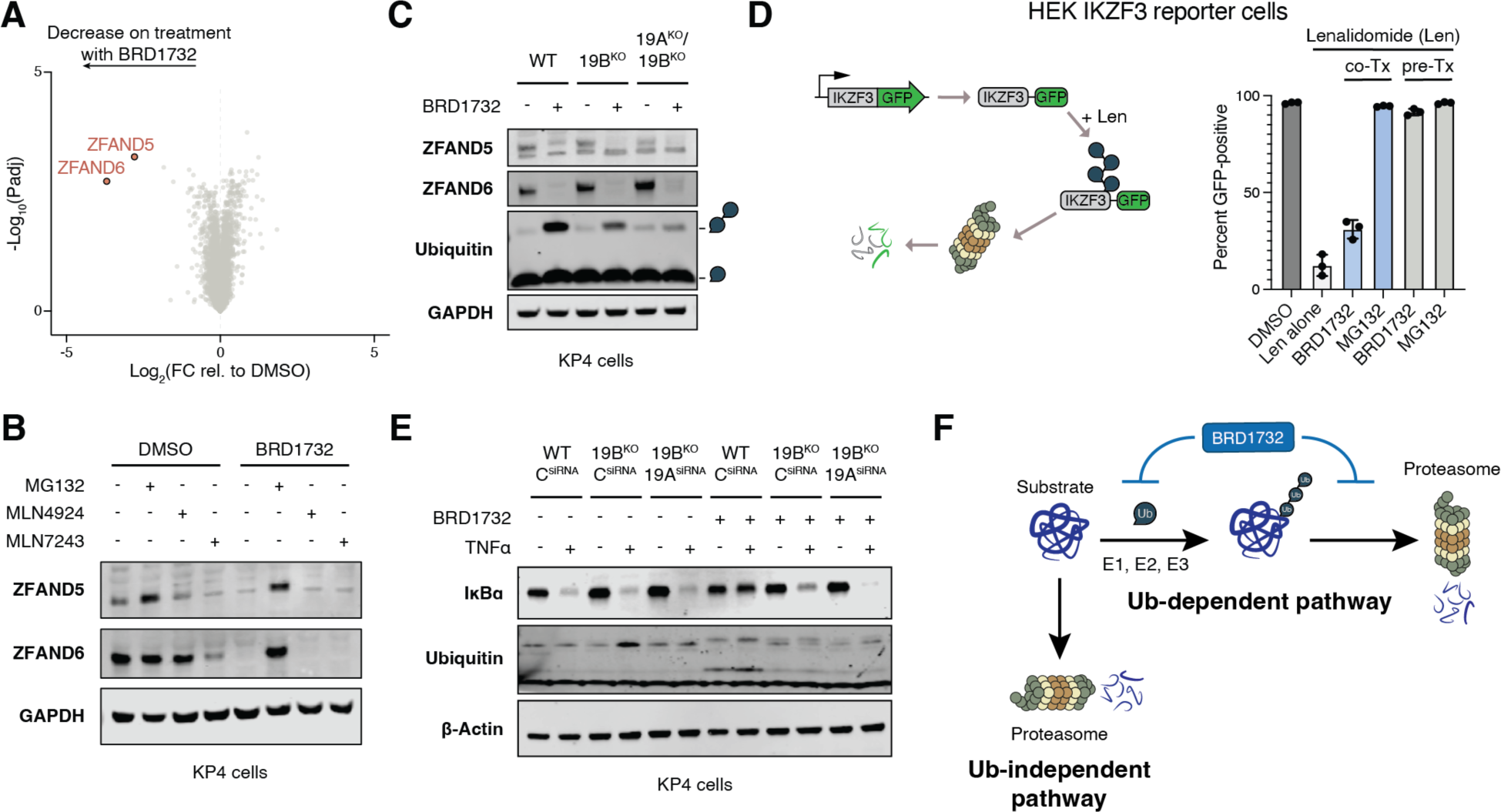
BRD1732 disrupts ubiquitin-dependent proteasomal degradation. (**A**) Quantitative proteomics from HCT116 cells treated with DMSO (n=3) or 5 µM BRD1732 (n=3) for 4 hours (two-tailed *t*-test). (**B**) ZFAND5 and ZFAND6 levels were analyzed by immunoblot from KP4 cells treated with 10 µM MG132, 500 nM MLN4924, or 50 nM MLN7243 in combination with either DMSO or 10 µM BRD1732 for 6 hours. (**C**) ZFAND5 and ZFAND6 levels were analyzed by immunoblot from KP4 wildtype, RNF19B knockout, or RNF19A/B double knockout cells following treatment for 6 hours with DMSO or 5 µM BRD1732. (**D**) Lenalidomide induces proteasomal degradation of IKZF3-GFP. HEK293T cells stably expressing IKZF3-GFP were subjected to treatment with DMSO, 5 µM BRD1732, or 5 µM MG132 alone or in combination with lenalidomide, and GFP levels were quantified by flow cytometry. Len, Lenalidomide; co-Tx, co-treatment for 2 hours; pre-Tx, pre-treatment with BRD1732 or MG132 for 6 hours prior to addition of lenalidomide for 2 hours. (**E**) KP4 wildtype or RNF19B knock out cells were transfected with RNF19A siRNA or control siRNA, subjected to 6-hour pre-treatment with BRD1732 or DMSO followed by 20-min stimulation with TNFα. IκBα degradation and polyubiquitin accumulation were analyzed by immunoblot. (**F**) Proposed model wherein BRD1732 blocks ubiquitin-dependent proteasomal degradation without inhibiting ubiquitin-independent proteasomal degradation.

Based on our finding that BRD1732 covalently modifies ubiquitin in cells and induces bidirectional effects on protein levels, we suspected that BRD1732 interferes with ubiquitin-dependent proteasomal degradation while leaving ubiquitin-independent degradation pathways intact. To characterize the effect of BRD1732 on ubiquitin-dependent degradation we evaluated its effect on lenalidomide-induced degradation of IKZF3 using a reporter system (*5*). Lenalidomide induces ubiquitination and degradation of IKZF3 by the E3 ubiquitin ligase cereblon, and this effect is blocked by co-treatment with MG132. While simultaneous co-treatment with BRD1732 and lenalidomide only minimally affected IKZF3 degradation, pre-treatment with BRD1732 for 6-hours robustly blocked degradation induced by lenalidomide (Fig. 4D).

To assess the effect of BRD1732 on ubiquitin-dependent proteasomal degradation in an endogenous system, we evaluated TNFα-induced proteasomal degradation of IκBα (Fig. 4E). Again with 6-hour pretreatment, BRD1732 efficiently blocked degradation of IκBα upon stimulation with TNFα. As expected, cells lacking expression of one or both RNF19 proteins showed reduced production of the ubiquitin-BRD1732 conjugate and impaired inhibition of ubiquitin-dependent proteasomal degradation. Our results are consistent with a model in which BRD1732 is ubiquitinated by RNF19 proteins, and accumulation of the resulting ubiquitin-BRD1732 conjugate leads to competitive inhibition of the ubiquitin-proteasome system (Fig. 4F).

## Discussion

Small molecule mechanism of action studies have repeatedly helped lay the groundwork for new fields of drug discovery. However, even when a target can be identified, achieving a deep understanding of a small molecule’s mechanism remains a significant challenge. Powerful modern tools such as genomic-wide CRISPR screening can be illuminating, but these are inadequate for unveiling the details of a chemical mechanism, which is still only achievable through highly-tailored biochemical studies.

Here, we use a combination of CRISPR screening, proteomics, and classical biochemistry to uncover a previously unknown small molecule mechanism of action wherein BRD1732 acts as a miniature protein to receive a posttranslational modification, ubiquitin. Nearly all known covalent small molecule probes and drugs contain reactive *electrophiles* to allow covalent modification of *nucleophilic* amino acids, such as cysteine, serine, or tyrosine, on their protein targets. Here, the reactive moiety resides within the small molecule. For BRD1732, the directionality of the reaction is reversed. BRD1732 contains a relatively unreactive *nucleophile*, the azetidine nitrogen, which forms a covalent bond with the C-terminus of ubiquitin. In this case, the reactive *electrophile* is the thioester bond between the ubiquitin C-terminus and the catalytic cysteine of the ubiquitin ligase.

Nucleotide-mimetic inhibitors of E1 enzymes also form covalent conjugates with the C-terminus of ubiquitin or NEDD8. However, while E1 enzyme inhibitors remain locked in the enzyme active site after conjugation, BRD1732 is ubiquitinated processively, and the resulting conjugate accumulates dramatically in cells to such an extent that free ubiquitin is reduced. Given the wide-ranging functions and interactors of ubiquitin in the cell, it is unsurprising that BRD1732 shows pleiotropic effects culminating in disruption of the ubiquitin-proteasome system and proteasomal degradation. While it shows similarity with active site proteasome inhibitors at the transcriptional and proteomic level, BRD1732 is unique in causing widespread bidirectional effects on protein homeostasis.

Ubiquitination of BRD1732 is both highly stereospecific and highly dependent on expression of RNF19 proteins. While we have been unable to reconstitute ubiquitination of BRD1732 by RNF19 with recombinant proteins despite multiple attempts, based on our CRISPR-Cas9 knockout and subsequent re-expression experiments, it is highly likely that RNF19 proteins are directly responsible for ubiquitination of BRD1732. Ubiquitination of non-proteinaceous substrates has only rarely been reported, notably with ubiquitination of lipopolysaccharide by RNF213 upon entry of *Salmonella* into the cytosol (*18*). Our demonstration of ubiquitination of a small molecule in cells raises the possibility of non-covalently directing ubiquitin to any protein target using bifunctional compounds related to BRD1732 within the cell. Indeed, our work more generally raises the possibility of posttranslational modification *in trans*, the delivery of a posttranslational mark through a small molecule intermediary.

---

**Fig. S1.**
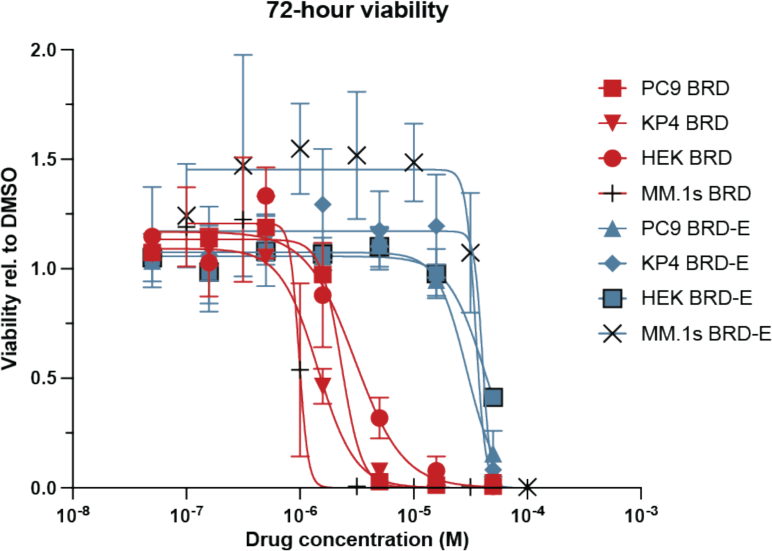
BRD1732 viability effects are stereospecific across a range of cell line lineages. Cell viability after 72-hour drug treatment of BRD1732 (BRD) and BRD-E for indicated cell lines. Fit was generated by 4-variable non-linear regression (n=3 technical replicates, n=3 biological replicates).

**Fig. S2.**
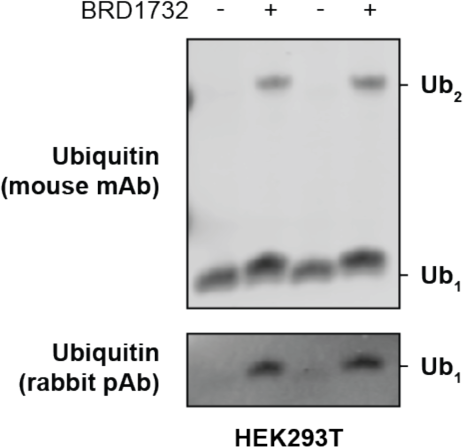
BRD1732 induces gel-shift for mono-ubiquitin. HEK293T cells were treated with DMSO or BRD1732 for 6 hours and ubiquitin levels were analyzed by immunoblot following separation on 12% polyacrylamide gel (representative of two independent experiments).

**Fig. S3.**
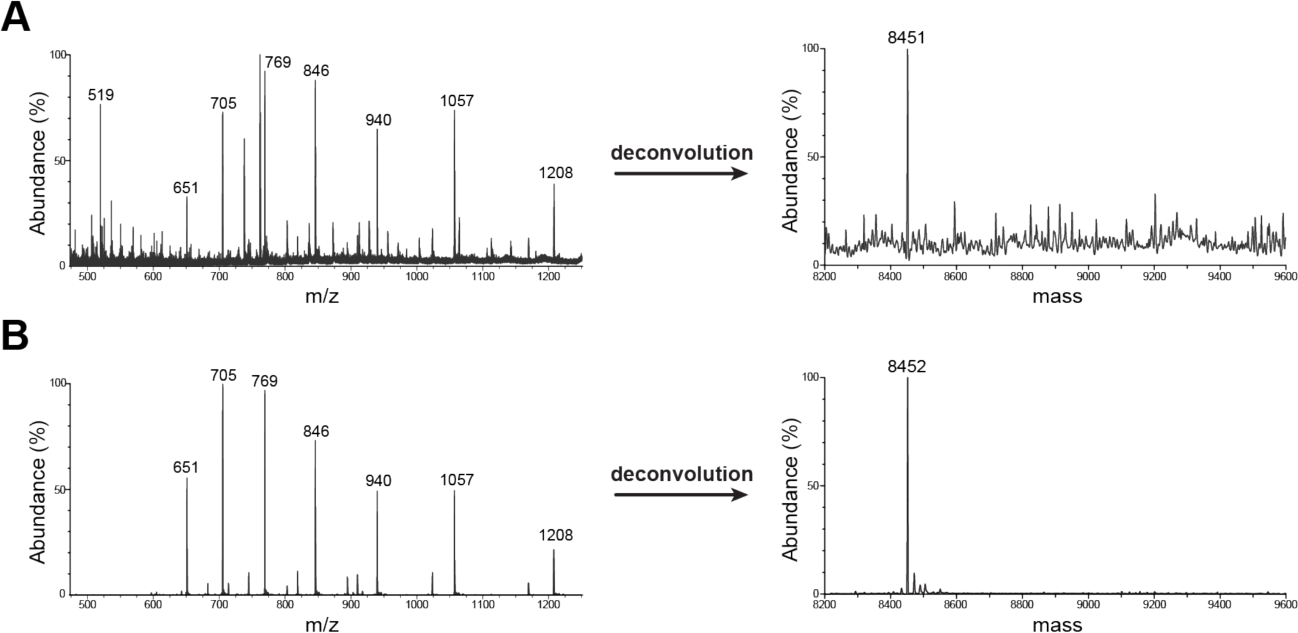
LC-MS of ubiquitin and ubiquitin conjugate following trypsin digestion. (**A**) Intact protein LC-MS of ubiquitin purified from BRD1732-treated Expi293F cells treated with trypsin under non-denaturing conditions (data also shown in Fig 2C). (**B**) Intact protein LC-MS of recombinant ubiquitin treated with trypsin under non-denaturing conditions.

**Fig. S4.**
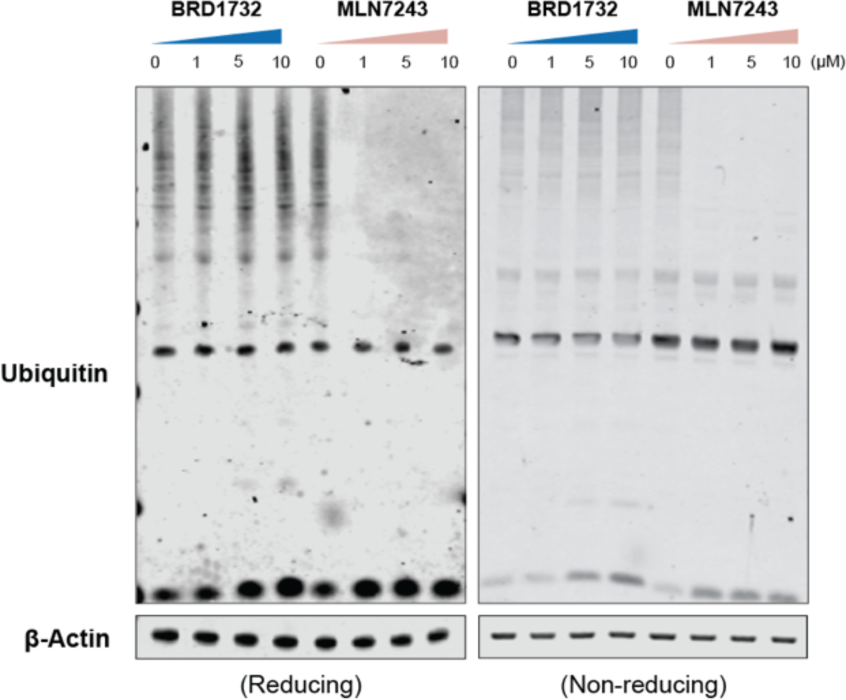
MLN7243 does not induce low-molecular weight ubiquitin polymers like BRD1732. Immunoblot following treatment of KP4 cells with indicated concentrations of BRD1732 or MLN7243 for 6 hours. Same experiment presented in Fig. 3C, now showing entire ubiquitin blot. Representative of two independent experiments.

## Methods

### Chemical Synthesis General Procedures

Oxygen and/or moisture sensitive reactions were carried out in oven or flame-dried glassware under nitrogen atmosphere. All reagents and solvents were purchased and used as received from commercial vendors or synthesized according to cited procedures. Yields refer to chromatographically and spectroscopically pure compounds, unless otherwise stated. Flash chromatography was performed using 20-40 µm silica gel (60 Å mesh) on a Teledyne Isco Combiflash Rf. Analytical thin layer chromatography (TLC) was performed on 0.25 mm silica gel 60-F plates and visualized by UV light (254 nm) and/or iodine vapor.

NMR spectra were recorded on Bruker 300 (^1^H, 300 MHz; ^13^C, 75 MHz) or 400 (^1^H, 400 MHz; ^13^C, 100 MHz) spectrometers. NMR solvents were purchased from Cambridge Isotope Laboratories, Inc., and NMR data were collected in CDCl_3_ at ambient temperature unless otherwise noted. Chemical shifts are reported in parts per million (δ, ppm) relative to CDCl_3_ (^1^H, 7.26; ^13^C, 77.16) or methanol-*d*_4_ (^1^H, 3.31; ^13^C, 49.00). Data for ^1^H NMR are reported as follows: chemical shift, multiplicity (br = broad, s = singlet, d = doublet, t = triplet, q = quintuplet, m = multiplet), coupling constants (J) in Hz, and integration. Purity was measured by LC-MS on a Waters 2795 separations module by UV absorbance at 210 nm and identity was determined on a SQ mass spectrometer by positive (M+H)^+^ or negative (M-H)^-^ electrospray ionization. Mobile phase A consisted of 0.01% formic acid in water, while mobile phase B consisted of 0.01% formic acid in acetonitrile. The gradient ran from 5% to 95% mobile phase B over 2.5, 5.0, or 7.5 minutes at 1.75 mL/min. An Agilent Poroshell 120 ECC18, 2.7 um, 3.0×30 mm column was used with column temperature maintained at 40°C.

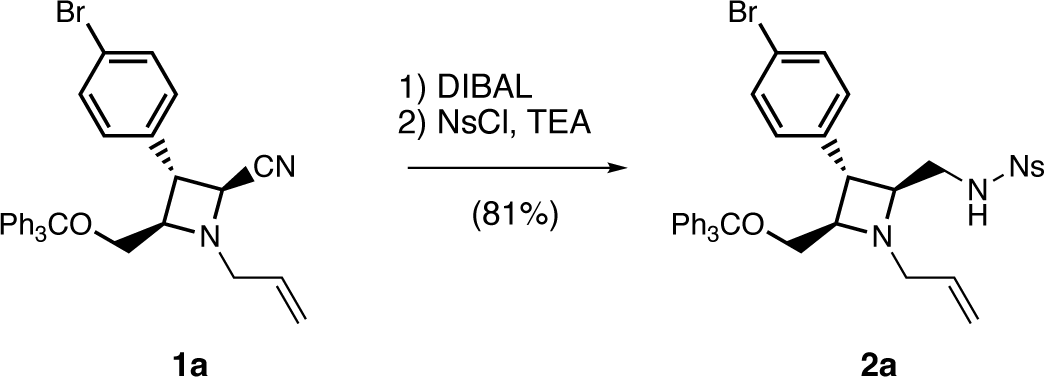

**N-(((2S,3R,4R)-1-allyl-3-(4-bromophenyl)-4-((trityloxy)methyl)azetidin-2-yl)methyl)-4-nitrobenzenesulfonamide (2a).** To a solution of (2S,3R,4R)-1-allyl-3-(4-bromophenyl)-4-((trityloxy)methyl)azetidine-2-carbonitrile (1a, as described in (*19*)) (1.0 g, 1.0 equiv) in DCM (18.2 mL, 0.1 M) at 0 °C was added neat DIBAL (1.95 mL, 6.0 equiv) dropwise over 20 min. The reaction was slowly warmed to ambient temperature (23 °C) over 2 h. Upon complete reduction, the reaction was cooled to 0 °C and MeOH was added dropwise (*note: vigorous bubbling and significant exotherm observed*) until bubbling ceased. Next, a solution of saturated Rochelle’s salt (potassium sodium tartrate in H_2_O, 15 mL per 1.0 mmol of substrate **1a**) was added dropwise over 5 min (*note: significant exotherm observed*), and the mixture was stirred at 0 °C for 2 h. The mixture was then extracted 4x with DCM, and combined organic layers were dried over Na_2_SO_4_, filtered, and removed *in vacuo* to afford crude primary amine as a thick yellow oil.

The resulting crude oil was reconstituted in DCM (18.2 mL, 0.1 M) and cooled to 0 °C before the addition of Et_3_N (634 µL, 2.5 equiv) and 2-nitrobenzenesulfonyl chloride (424 mg, 1.05 equiv). After 15 min or complete conversion, the solvent was removed *in vacuo* and crude residue purified using flash column chromatography (EtOAc/hexanes) to afford desired product as a white foaming solid (1.09 g, 81%). ^1^H NMR (400 MHz, CDCl_3_) δ 8.12 – 8.05 (m, 1H), 7.84 (dd, J = 6.9, 2.2 Hz, 1H), 7.78 – 7.68 (m, 2H), 7.45 – 7.39 (m, 7H), 7.35 – 7.23 (m, 10H), 7.01 (d, J = 8.0 Hz, 2H), 6.12 (s, 1H), 5.67 (dq, J = 16.8, 8.4, 8.0 Hz, 1H), 5.11 (d, J = 17.2 Hz, 1H), 4.85 (d, J = 10.1 Hz, 1H), 3.38 (ddd, J = 28.3, 11.3, 4.9 Hz, 2H), 3.23 (q, J = 9.3, 7.1 Hz, 5H), 3.14 – 2.98 (m, 2H). Molecular formula: C_39_H_36_BrN_3_O_5_S. Calculated mass: 737.16. ESIMS m/z 738.4 (M+H)^+^.

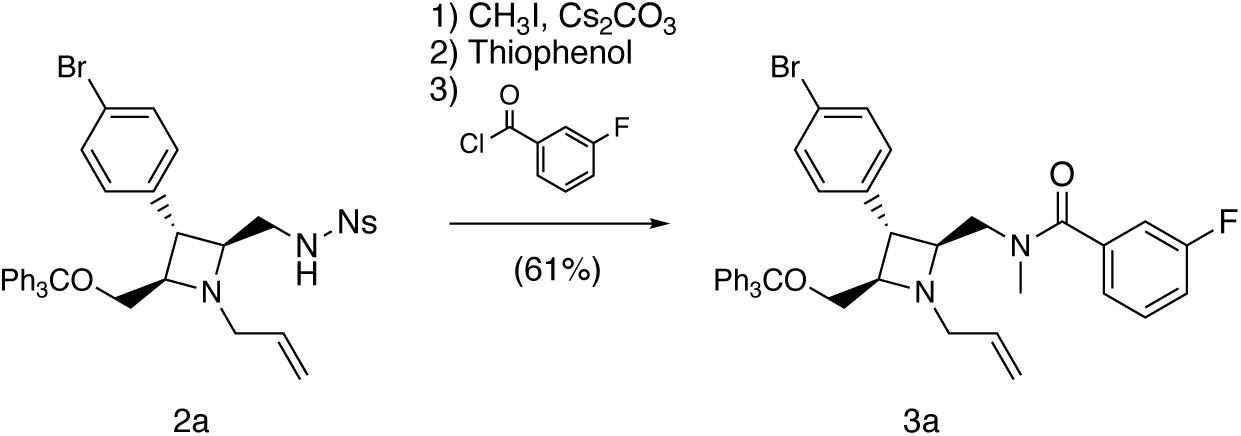

**N-(((2S,3R,4R)-1-allyl-3-(4-bromophenyl)-4-((trityloxy)methyl)azetidin-2-yl)methyl)-3-fluoro-N-methylbenzamide (3a).** To a solution of **2a** (300 mg, 1.0 equiv) in DMF (4 mL, 0.1 M) was added Cs_2_CO_3_ (198 mg, 1.5 equiv) followed by methyl iodide (28 µL, 1.1 equiv) and allowed to stir for 16 h at ambient temperature. The reaction was then quenched by the addition of water and extracted 3x with Et_2_O. The combined organic layers were dried over Na_2_SO_4_, filtered, and removed *in vacuo* to afford a thick crude yellow residue.

The residue was reconstituted in DMF (4 mL, 0.1 M), followed by the addition of K_2_CO_3_ (337 mg, 6 equiv) and thiophenol (133 µL, 3 equiv). The reaction was allowed to stir at ambient temperature or until completion, at which point saturated NaHCO_3_ was added to quench the reaction. The aqueous layer was extracted 3x with EtOAc, and combined organic layers were dried over Na_2_SO_4_, filtered, and removed *in vacuo*. The crude material was loaded onto a short silica column and quickly washed with DCM to elute excess thiophenol. Approximately 4 column volumes of 20% MeOH/DCM was then added to elute the diamine 1-((2S,3R,4R)-1-allyl-3-(4-bromophenyl)-4-((trityloxy)methyl)azetidin-2-yl)-N-methylmethanamine.

The diamine was dissolved in DCM (4 mL, 0.1 M) and cooled to 0 °C. Et_3_N (170 µL, 3.0 equiv) and 3-fluorobenzoyl chloride (57 µL, 1.15 equiv) were added in sequence. After 15 min, the reaction was quenched with saturated NH_4_Cl and extracted 3x with DCM. The combined organic layers were dried over Na_2_SO_4_, filtered, and removed *in vacuo*, and crude material was purified using flash column chromatography (MeOH/DCM) to afford desired product as a white foaming solid (170 mg, 61% over three steps). ^1^H NMR (400 MHz, CDCl_3_) δ 7.52 – 7.38 (m, 8H), 7.37 – 7.22 (m, 12H), 7.14 – 6.86 (m, 3H), 6.05 – 5.68 (m, 1H), 5.31 (d, J = 16.3 Hz, 1H), 5.20 – 4.99 (m, 1H), 3.95 (dd, J = 14.0, 3.7 Hz, 1H), 3.62 – 3.11 (m, 8H), 3.04 (d, J = 28.4 Hz, 3H). Molecular formula: C_41_H_38_BrFN_2_O_2_. Calculated mass: 688.21. ESIMS m/z 689.1 (M+H)^+^.

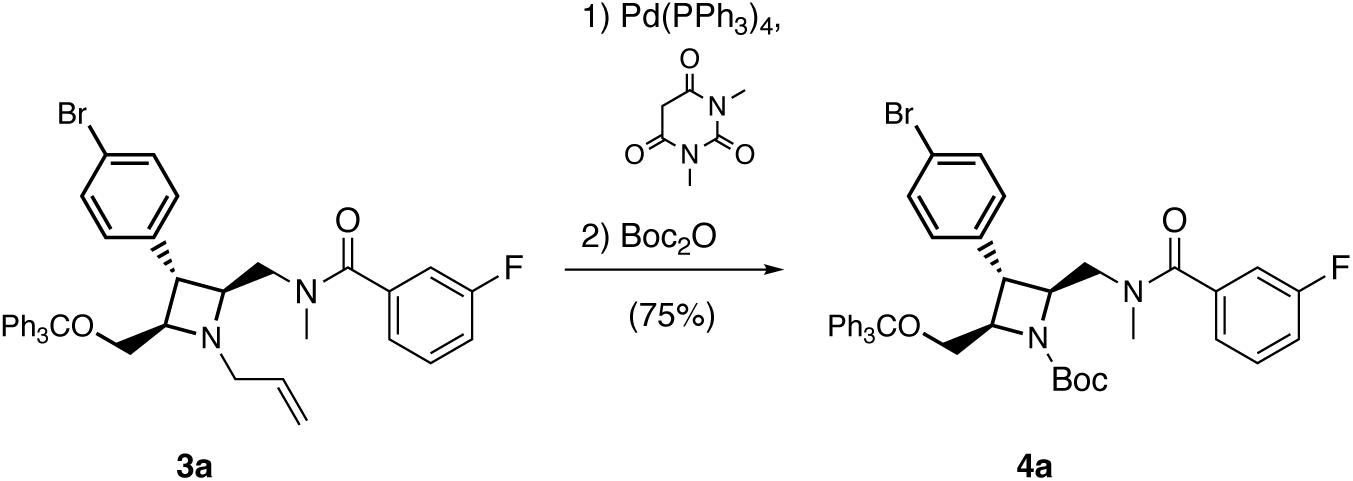

**tert-butyl (2S,3R,4R)-3-(4-bromophenyl)-2-((3-fluoro-N-methylbenzamido)methyl)-4-((trityloxy)methyl)azetidine-1-carboxylate (4a)**. To a solution of **3a** (170 mg, 1.0 equiv) in 2:1 EtOH/DCM (4.9 mL, 0.05 M) was added 1,3-dimethylbarbituric acid (58 mg, 1.5 equiv) and Pd(PPh_3_)_4_ (29 mg, 0.1 equiv). The mixture was heated to 40 °C for 4 h, or until complete deallylation was observed, at which point Boc_2_O (65 mg, 1.2 equiv) was added in a single portion. After 1 h at 40 °C, the solvent was removed *in vacuo* and crude material was purified using flash column chromatography (EtOAc/hexanes) to afford desired product as a pale yellow foaming solid (139 mg, 75%). ^1^H NMR (400 MHz, CDCl_3_) δ 7.60 – 7.18 (m, 19H), 7.17 – 6.90 (m, 4H), 4.50 – 4.35 (m, 1H), 4.25 (s, 1H), 4.09 (d, J = 16.1 Hz, 1H), 3.98 – 3.82 (m, 1H), 3.65 (t, J = 6.9 Hz, 1H), 3.48 (d, J = 4.9 Hz, 2H), 3.33 (s, 1H), 3.01 (s, 3H), 1.45 (s, 9H). Molecular formula: C_43_H_42_BrFN_2_O_4_. Calculated mass: 749.28. ESIMS m/z 773.3 (M+H+Na^+^)^2+^.

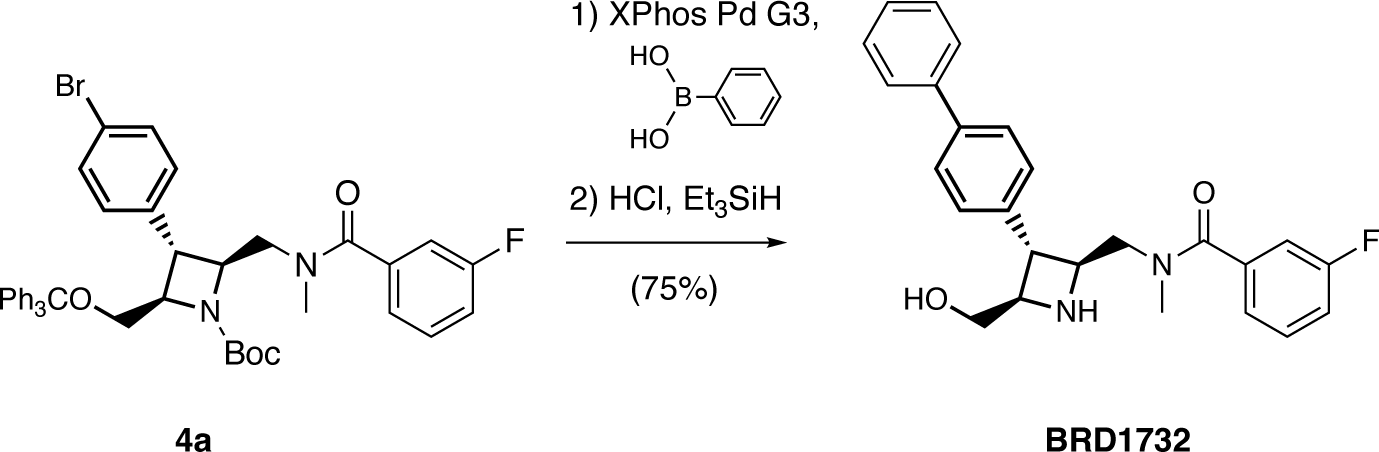

**N-(((2S,3R,4R)-3-([1,1’-biphenyl]-4-yl)-4-(hydroxymethyl)azetidin-2-yl)methyl)-3-fluoro-N-methylbenzamide (BRD1732).** A reaction vessel charged with **4a** (25 mg, 1.0 equiv) and phenylboronic acid (8.1 mg, 2.0 equiv) was evacuated and backfilled 3x with N_2_. Degassed solutions of dioxane (333 µL, 0.1 M) and 0.5 M K_3_PO_4_ (200 µL, 3.0 equiv) were added, followed by XPhos Pd G3 (4.2 mg, 0.15 equiv), and the mixture was heated to 100 °C. After 1 h or upon complete conversion, the mixture was quenched with saturated NH_4_Cl and extracted 3x with EtOAc. The combined organic layers were dried over Na_2_SO_4_, filtered, and removed *in vacuo* to afford a dark brown residue.

The residue was reconstituted in 4 M HCl in dioxane (830 µL, 100 equiv), followed by the addition of Et_3_SiH (8 µL, 1.5 equiv). After 1 h, solvent was removed *in vacuo* and crude material was purified using flash column chromatography (MeOH/DCM) to afford desired product as a pale yellow residue (7.5 mg, 53%). ^1^H NMR (400 MHz, MeOD) δ 7.65 – 7.59 (m, 4H), 7.49 (d, *J* = 7.8 Hz, 2H), 7.46 – 7.41 (m, 3H), 7.38 – 7.32 (m, 2H), 7.19 – 7.12 (m, 2H), 6.95 (d, *J* = 7.6 Hz, 1H), 6.89 (d, *J* = 9.0 Hz, 1H), 4.09 (dd, *J* = 8.7, 4.4 Hz, 1H), 4.00 (dd, *J* = 13.7, 6.7 Hz, 1H), 3.78 – 3.62 (m, 4H), 3.55 (s, 1H), 2.95 (s, 3H). Molecular formula: C_25_H_25_FN_2_O_2_. Calculated mass: 404.19. ESIMS m/z 405.3 (M+H)^+^.

**N-(((2R,3S,4S)-3-([1,1’-biphenyl]-4-yl)-4-(hydroxymethyl)azetidin-2-yl)methyl)-3-fluoro-N-methylbenzamide (BRD-E).** The enantiomer of BRD1732 was synthesized from (2R,3S,4S)-1-allyl-3-(4-bromophenyl)-4-((trityloxy)methyl)azetidine-2-carbonitrile (prepared as in (*19*)) in a manner analogous to BRD1732. ^1^H NMR (400 MHz, CD_3_OD) δ 7.65 – 7.59 (m, 4H), 7.49 (d, J = 7.8 Hz, 2H), 7.46 – 7.41 (m, 3H), 7.38 – 7.32 (m, 2H), 7.19 – 7.12 (m, 2H), 6.95 (d, J = 7.6 Hz, 1H), 6.89 (d, J = 9.0 Hz, 1H), 4.09 (dd, J = 8.7, 4.4 Hz, 1H), 4.00 (dd, J = 13.7, 6.7 Hz, 1H), 3.78 – 3.62 (m, 4H), 3.55 (s, 1H), 2.95 (s, 3H). Molecular formula: C_25_H_25_FN_2_O_2_. Calculated mass: 404.19. ESIMS m/z 405.0 (M+H)^+^.

**N-(((2R,3R,4S)-3-([1,1’-biphenyl]-4-yl)-4-(hydroxymethyl)azetidin-2-yl)methyl)-3-fluoro-N-methylbenzamide (BRD-D).** The (2R,3R,4S) diastereomer of BRD1732 was synthesized from (2R,3R,4S)-1-allyl-3-(4-bromophenyl)-4-((trityloxy)methyl)azetidine-2-carbonitrile (prepared as in (*19*)) in a manner analogous to BRD1732. ^1^H NMR (400 MHz, DMSO d6) δ 8.30 (s, 1H), 7.75 – 7.64 (m, 2H), 7.63 – 7.57 (m, 2H), 7.55 – 7.51 (m, 4H), 7.50 – 7.24 (m, 3H), 7.22 – 7.12 (m, 2H), 4.36 (m, 1H), 4.05 (m, 1H), 3.81 – 3.38 (m, 4H), 3.22 – 3.09 (m, 1H), 2.88 (s, 1.5H), 2.74 (s, 1.5H). Molecular formula: C_25_H_25_FN_2_O_2_. Calculated mass: 404.19. ESIMS m/z 405.5 (M+H)^+^.

### Antibodies and expression constructs

The details and sources of the antibodies and plasmid constructs that were used in this study are summarized in the tables at the end of this section. In addition, human RNF19A variant constructs were synthesized and cloned into the pLVX6 backbone by Twist Biosciences. All constructs were verified by whole-plasmid sequencing (Primordium Labs).

### Cell culture and reagents

PC9, MM.1S, and HEK293T cells were purchased from ATCC. HCT116 cells were obtained from the Broad Institute Genetic Perturbation Platform. KP4 cells were purchased from JCRB Cell Bank (Japan). HEK293T IKZF3 reporter cells were a gift from the laboratory of Benjamin Ebert (DFCI). Expi293F cells were purchased from Thermo Scientific. All cell lines were confirmed to be mycoplasma free. Transient transfections were carried using TransIT-LT1 reagent (Mirus Bio). HEK293T, HCT116 and KP4 cells were maintained in DMEM medium (Gibco) containing 10% (v/v) fetal bovine serum (FBS) (Gibco). PC9 and MM.1S cells were maintained in RPMI medium (Gibco) containing 10% FBS (Gibco). Expi293F cells were maintained in Expi293 Expression Medium (Thermo Scientific).

### Cell viability assays

Cells were seeded in 96-well plates at 1,000 cells per well in 90 μl DMEM with 10% FBS. Following attachment for 24 hours, cells were treated by addition of 10 μl compound dilution or vehicle (0.1% DMSO final). After 72 hours, cell viability was quantified using the CellTiterGlo Luminescent Cell Viability Assay (Promega) measuring luminescence on a SpectraMax M3 plate reader or BioTek Synergy 2 plate reader. Data were processed using GraphPad Prism.

### Multiplexed quantitative proteomic analysis in Yamato cells

Yamato synovial sarcoma cells were plated in 10cm dishes at a density of 2×10^6^ cells / mL and allowed to grow for 48 hours or until 75% confluency was reached. These cells were treated with BRD1732 at 1 or 5 μM, MG132 at 5 μM, or DMSO for 24 hours at a final DMSO concentration of 1% v/v. Cells were harvested, washed with ice-cold PBS, and lysed in 200μL of RIPA (50 mM Tris-HCl pH 7.5, 150 mM NaCl, 1% Triton X-100, 0.1% SDS, 0.5% sodium deoxycholate, 1mM DTT, and cOmplete Protease Inhibitors (Roche)). Protein concentrations were measured using the DC Protein Assay (Bio-Rad). Data were acquired and analyzed as previously described (*20*). Adjusted p-value was calculated by student’s t-test followed by Benjamini-Hochberg procedure. Proteomic data sets were processed to remove proteins with expression changes by RNA-Seq (|Log_2_FC|>1 and -Log_10_(*p*_adj_)>2, where *p*_adj_ is BH-corrected *p*-value). Resulting data sets for 5 µM BRD1732 and 5 µM MG132 were compared to identify proteins with Log_2_FC>1.5 and - Log_10_(*p*_adj_)>2 (where *p*_adj_ is BH-corrected p-value). Plots were generated with GraphPad Prism.

### RNA-Seq analysis in Yamato cells

Yamato cells were plated in 15cm dishes at a density of 2×106 cells / mL and allowed to grow for 48 hours or until 75% confluency was reached. These cells were then treated with BRD1732 at 5µM, MG132 at 5µM, or DMSO in triplicate at a final DMSO concentration of 1% v/v, and incubated for 24 hours. Following perturbation, media was removed and 1mL of TRIzol reagent (ThermoFisher) was added to each well, and RNA was extracted according to the manufacturers protocol. Quantification of the samples was performed using the Qubit RNA HS Assay Kit (ThermoFisher) in a Qubit fluorometer (ThermoFisher). All library preparation and sequencing (75bp single end on Illumina NextSeq 500) was performed in the Molecular Biology Core Facilities at the Dana-Farber Cancer Institute. RNA-Seq reads were mapped to the human reference genome (hg19) using STAR version 2.3.1 (*21*) with default parameters. All error bars represent Mean±SEM. Significance was assessed using the R package DESeq2 (*22*) using raw read counts generated with Rsubread featureCounts against the hg19 refFlat annotation.

### Genome-wide CRISPR resistance screening

KP4 cells expressing Cas9 were infected with the AVANA genome-wide sgRNA library (*23*) and plated in T175 flasks at a density of 2×106 cells / mL and allowed to adhere for 48 hours. Cells were then treated with BRD1732 at 10μM or DMSO for 3 days, and allowed to recover from treatment for additional 7 days. Following recovery cells were harvested and genomic DNA was isolated using Maxi kits according to the manufacturer’s protocol (Qiagen). Next-generation sequencing was performed by the Genomics Platform at the Broad Institute.

MM.1S multiple myeloma cells engineered to stably express the nuclease SpCas9 (provided by Quinlan L. Sievers, B. Ebert lab) were also transduced with a pool of lentiviral particles for ∼70,000 sgRNAs (Brunello library), targeting exons of ∼20,000 genes (plus non-targeting control sgRNAs), under conditions of transduction which allow for an average of no more than 1 sgRNA to be incorporated in a given cell (*24*). The MM.1S-Cas9+ Brunello-transduced cells were then cultured in the presence or absence of increasing doses of BRD1732 (0.5-4 µM). The treated vs. control cultures of MM.1S cells were passaged serially over 10 weeks and were periodically checked for BRD-1732 sensitivity using CTG. When the dose-responses curve identified the isolation of BRD-1732 resistant MM cells, these were isolated from the cultures and processed for sequencing. Data was analyzed with the MAGeCK package using default parameters.

### Multiplexed quantitative proteomics time-course and DeepCoverMOA

HCT116 cells were plated in 6-well plates at a density of 1×10^6^ cell/well. After attachment for 24 hours, cells were treated with 5 µM BRD1732 for 1, 2, 4, or 24 hours or 0.5% (v/v) DMSO for 24 hours. Cells were then washed with PBS three times and lysed with 8M Urea, 0.1% SDS, 200mM EPPS buffer pH 8.5, containing cOmplete Protease Inhibitor Cocktail Tablets (Roche) and PhosSTOP (Roche). Samples were processed as previously described (*17*) but using 16-plex TMTpro reagents. Briefly, 30 μg of protein was reduced with TCEP (5mM, 15min), alkylated with iodoacetamide (20mM, 30min) then extracted using single-pot, solid-phase-enhanced sample-preparation (SP3) technology. Proteins were digested overnight at 37C using 0.3μg LysC (Wako) and 0.3μg Trypsin (Thermo) before TMT labeling. TMT labeled peptides were pooled then desalted using a 50mg SepPak (Waters) before offline basic pH reversed-phase HPLC fractionation, generating 24 fractions. 12 non-adjacent fractions were desalted using in-housed packed stagetips, then analyzed using a real-time search MS3 method. Adjusted *p*-value was calculated by student’s t-test followed by Benjamini-Hochberg procedure. Plots were generated using GraphPad Prism. Data corresponding to Log_2_FC from 24-hour time point were subjected to comparative analysis by DeepCoverMOA (http://wren.hms.harvard.edu/DeepCoverMOA/) (*17*). When limiting analysis to proteins that increase upon treatment with BRD1732, data for proteins with Log_2_FC<0 were removed and remaining data were subjected to analysis by DeepCoverMOA. Networks were generated using Cytoscape.

### Immunoblotting

Cells were plated into 12-well plate (Costar, #3513) with 100,000 cells per well (final volume 1 mL) and incubated overnight. Cells at 40-60% confluency were treated with compounds or 0.5% (v/v) DMSO. After drug treatment for indicated times, cells were lysed in CelLytic M (Sigma-Aldrich) reagent supplemented with 1 x EDTA-free protease inhibitor cocktail tablet (Sigma-Aldrich) for 40 min on a rotating wheel at 4°C. Lysates were separated on 4-12% Bis-Tris SDS PAGE (Invitrogen) in MES buffer (Invitrogen) at 150V for 45 min. Proteins were transferred to PVDF membrane using dry transfer method via iBlot 2 (Invitrogen) following the manufacturer’s instructions. The membrane was blocked with 5% BSA (RPI) in TBS-T buffer at room temperature for 1 hour followed by incubation with primary antibody at 4°C overnight, followed by secondary antibody incubation for 1 hour at room temperature. Immunoblots were then imaged by Li-Cor Odyssey CLx Imaging System. All antibodies are listed at the end of the section of materials and methods.

### PRISM profiling

PRISM profiling was performed as previously described (*25*). The PRISM cell set used consisted of 483 solid tumor cancer cell lines. These cell lines largely overlap with and reflect the diversity of the Cancer Cell Line Encyclopedia (CCLE) cell lines (see https://portals.broadinstitute.org/ccle). Cell lines were grown in RPMI 10% FBS without phenol red. Parental cell lines were stably infected with a unique 24-nucleotide DNA barcode via lentiviral transduction and blasticidin selection. After selection, barcoded cell lines were expanded and QCed (mycoplasma contamination test, a SNP test for confirming cell line identity, and barcode ID confirmation). Passing barcoded lines were then pooled (20-25 cell lines per pool) based on doubling time and frozen in assay-ready vials. A Labcyte Echo was used to acoustically transfer test compounds were to 384-well plates at 8 doses with 3-fold dilutions in triplicate. These assay ready plates were then seeded with the thawed cell line pools. Adherent cell pools were plated at 1250 cells per well. Treated cells were incubated for 5 days then lysed with QIagne TCL buffer and frozen. Lysate plates were collapsed together prior to barcode amplification and detection. Each cell line’s unique barcode is located at the end of the blasticidin resistance gene and gets expressed as mRNA. These mRNAs were then captured by using magnetic particles that recognize polyA sequences. mRNA was then reverse-transcribed into cDNA and then the sequence containing the unique PRISM barcode was amplified using PCR. Finally, Luminex beads that recognize the specific barcode sequences in the cell set were hybridized to the PCR products and then detected using a Luminex scanner which reports signal as a median fluorescent intensity (MFI).

### Endogenous ubiquitin purification

Ubiquitin-BRD1732 conjugate was prepared by following steps. Expi293T cells were grown in in 125-mL shaker flasks at 37°C and 8% CO_2_. Cells were transiently transfected with pLVX6 RNF19A using the ExpiFectamine 293 Transfection Kit following the manufacturer’s instructions. After incubation overnight, media was exchanged to Expi293 Expression Media containing 10 µM BRD1732 and cells were incubated for 6 hours. The cell stock was then resuspended with cell lysis reagent CelLytic M (Sigma-Aldrich) supplemented with EDTA-free protease inhibitor cocktail tablet (Sigma-Aldrich) and incubated for 40 min at 4°C on a shaker. The sample were then centrifuged at 1,200 x g for 2 min using table centrifuge. The supernatant was collected and filtered using 0.45 µm filter (Millex) before loading on 1 mL HiTrap Q HP column (Cytiva) using fast protein liquid chromatography (FPLC) on ÄKTA pure (Cytiva). Ubiquitin-BRD1732 conjugate were eluted with a salt gradient from 0 to 1 M NaCl in 50 mM ammonium chloride, pH 6.0. The pooled protein sample was further purified by size-exclusion chromatography (SEC) using a Superdex 75 Increase 10/300 GL (Cytiva) column in PBS buffer. In comparison, wild-type ubiquitin was purified from Expi293T cells using the same mentioned-above method except the steps of pLVX6 plasmid transfection and BRD1732-treatment.

### Protein expression and purification

HA-tUI protein was purified as previously described (*11*). Briefly, HA-tUI protein cloned on pET28 (addgene #122662) were transformed into BL21 (DE3) Star II *Escherishia coli* (QB3-Berkeley) for protein expression. The expression was induced by adding 0.4 mM IPTG to cells grown at 37°C till OD_600_ reached 0.6 to 0.8. After a 30 min ice bath, the cells were continued to grow at 20°C overnight. The cells were then harvest at 6,000 x g for 25 min at 4°C. The collected pellets were resuspended in the Buffer A (20 mM sodium phosphate, 500 mM NaCl, 20 mM imidazole, 1 mM TCEP, pH = 7.4) and sonicated for cell lysis. The lysate is further centrifuged at 20,199 x g for 30 min at 4°C using JA-25.50 rotor. The supernatant was filtered against 0.45 µm filter (Millex) before loading on a 5 mL HisTrap HP column (Cytiva) for FPLC. The protein was eluted by the imidazole gradient from 20 mM to 500 mM in Buffer A. After an overnight buffer dialysis into buffer B (PBS, 1 mM TCEP, pH=7.4) where the protein solution was held in the 3.5 kDa cutoff tubing (Spectrum Labs), protein was then purified by SEC using a HiLoad 16/600 Superdex 200 pg (Cytiva) column in buffer B.

### Intact protein LC-MS

LC-MS were performed on a Xevo^®^ G2-XS QTOF mass spectrometer coupled with ACQUITY ultra performance liquid chromatography (UPLC) system equipped with ACQUITY UPLC protein BEH C4 column (300 Å, 1.7 µm, 2.1 mm x 50 mm). Buffer A water supplemented with 0.1% formic acid and buffer B acetonitrile supplemented with 0.1% formic acid were used as mobile phase. The flow rate is 0.5 mL/min. Spectra were deconvoluted and analyzed using Masslynx V4.2 software.

### Trypsin digestion and tandem MS-MS analysis of ubiquitin-BRD1732

FPLC-SEC purified ubiquitin-BRD1732 conjugate in PBS buffer (pH = 7.4) was trypsin-digested and analyzed by tandem MS. Briefly, purified ubiquitin-BRD1732 conjugate (∼1 µM, 100 µL) in PBS was buffer exchanged into digestion buffer A (20 mM Tris, pH = 8.0, 2 mM CaCl_2_) by using 0.5 mL 3 kDa-cutoff centrifugal filters (Amicon). Trypsin-digestion was performed by adding 0.5 mg trypsin into the sample and incubating overnight at 37°C. The digestion reaction was stopped by adding 5 µL 10% formic acid to reach a final concentration of 0.5 % (v/v) formic acid. Digested peptides were characterized by a Waters Xevo G2-XS system equipped with an Acquity UPLC BEH C4 1.7 µm column using a linear gradient of 5–95% acetonitrile/water + 0.05% formic acid. Gating on the ion at m/z=519 we applied collision energy of 20 eV. A similar protocol for trypsin digestion and intact protein LC-MS was utilized for unmodified ubiquitin.

### Quantification of free ubiquitin

HEK293T cells were plated 200, 000 cells per well in a 12-well plate and incubated at 37°C with 5% CO_2_ overnight. Drug treatments were performed by treating the cells with different concentration of BRD1732 for 6 hours. Cells were detached by trypsinization using TrypLE express enzyme (Thermo Fisher) and then washed by ice-cold PBS buffer (all the following PBS wash used ice-cold PBS buffer). Next, cells were fixed by incubating with 4% formaldyhyde (Sigma-Aldrich) in PBS buffer for 15 min at room temperature, followed by a PBS wash. Cells were then permeabilized by slowly dropping ice-cold 100% methanol (Fisher Scientific) with vertexing to the final concentration of 90% methanol, followed by 30 min incubation on ice and a PBS wash. Next, cell samples were incubated with blocking buffer (5% BSA (RPI) in TBST buffer) for 1 hour to block the non-specific binding. After that, samples were divided into two groups. Experimental group was stained with 100 nM purified HA-tUI protein in blocking buffer for 30 min at room temperature to detect the free ubiquitin concentration. In control group, purified 100 nM HA-tUI protein were mixing with 10 µM ubiquitin protein (Enzo life sciences) for 5 min at room temperature in blocking buffer before adding to the sample. Three PBS washes were performed with at least 5 min interval per each. Cells were then incubated overnight at 4°C with anti-HA mAb conjugated with Alexa Fluor 750 (Cell signaling) in the black box with 1:500 dilution. Next day, samples were washed three times with PBS buffer before loaded with 200 µL PBS supplemented with 1 % BSA. Samples were then transferred to 96-well plate and analyzed on a CytoFLEX S Flow Cytometer (Beckman). The results were then processed using FlowJo.

### IKZF3 reporter assay

HEK293T IKZF3 cells were plated at 3×10^5^ cells/well in 12-well plates. After attachment for 24 hours, cells were treated with 5 µM BRD1732, 5 µM MG132 or 0.1% (v/v) DMSO for 6 hours. Media was then exchanged to include 200 nM lenalidomide with indicated compounds for an additional 2 hours. Cells were detached using TrypLE Express (Thermo), resuspended in PBS with 10% (v/v) FBS, centrifuged and resuspended in PBS with 1% (v/v) FBS and analyzed by flow cytometry on a CytoFLEX S Flow Cytometer. Data were processed using FlowJo. Data represent three biological replicates.

### Generation of knock-out and double knock-out stable cells

KP4 or HEK293T cells were reverse transfected with preformed Cas9 RNP with EnGen Cas9 NLS (NEB), crRNA targeting RNF19B (target sequence: TCGTACTTGTGCATAAGCGG) and tracrRNA (IDT) using Lipofectamine CRISPRMAX (Thermo). Cells were allowed to attach for 24 hours then were selected with 5 µM BRD1732 for 24 hours to obtain RNF19B knockout (BKO) cells. Following two passages, BKO cells were similarly reverse transfected with Cas9 RNP with crRNA targeting RNF19A (target sequence: AAGAATTTATGCTTAGACGG). Following attachment for 24 hours, cells were selected with 20 µM BRD1732 for 24 hours to obtain RNF19A/B double knockout (DKO) cells.

### siRNA mediated knock-down

KP4 wildtype or BKO cells were reverse transfected with Silencer siRNA targeting RNF19A (134494, Thermo) or Silencer Select siRNA Control (AM4611, Thermo), plating cells at 8×10^5^ per well. Following attachment for 24 hours, cells were treated with 5 µM BRD1732 or DMSO 0.5% (v/v) DMSO for 6 hours followed by TNFα 20 ng/mL or water in media for 10 minutes. Plates were placed on ice and cells were harvested for immunoblot analysis by addition of cold RIPA buffer.

### Antibodies

| Designation | Catalog number | Source | Dilution |
| --- | --- | --- | --- |
| anti-Ubiquitin, Mouse IgG | 13-1600 | Invitrogen | 1:1000 |
| anti-Ubiquitin, Rabbit IgG | 3933S | Cell signaling | 1:1000 |
| anti-β-actin | 4970S/3700S | Cell signaling | 1:1000 |
| anti-HA (Alexa Fluor<sup>®</sup> 750) | 20818S | Cell signaling | 1:500 |
| anti-Ub-H2A-K119 | 8240S | Cell signaling | 1:500 |
| anti-E2T | 12992S | Cell signaling | 1:1000 |
| anti-E2A/B | 4944S | Cell signaling | 1:1000 |
| anti-Mcl-1 | 94296S | Cell signaling | 1:1000 |
| anti-Nrf1 | 8052S | Cell signaling | 1:1000 |
| anti-p21 | 2947S | Cell signaling | 1:1000 |
| anti-p27 | 3686S | Cell signaling | 1:1000 |
| anti-ZFAND5 | PA5-53735 | Invitrogen | 1:1000 |
| anti-ZFAND6 | PA5-57083 | Invitrogen | 1:1000 |
| anti-GAPDH | 97166S | Cell signaling | 1:1000 |
| anti-IκBα | 2859T | Cell signaling | 1:1000 |
| Goat anti-Mouse IgG<br>IRDye<sup>®</sup> 800CW | 926-68070 | LICOR | 1:10000 |
| Goat anti-Rabbit IgG<br>IRDye<sup>®</sup> 680RD | 926-32211 | LICOR | 1:10000 |

### Plasmids Sources

| Plasmid | Source |
| --- | --- |
| eGFP, pLVX6 | Gift from Zhenkun Lou |
| Human RNF19A, pLVX6 | Gift from Zhenkun Lou |
| Human RNF19B, pLVX6 | This study |
| Human RNF19A C316K, pLVX6 | This study |
| Human RNF19A RBR2TM, pLVX6 | This study |
| Human RNF19A RBR1TM, pLVX6 | This study |
| Human RNF19A RBR, pLVX6 | This study |
| Human Ubb<sup>+1</sup> with myc tag, pCMV-Myc | Gift from Nico Dantuma<br>Addgene plasmid # 20254 |
| Human ubiquitin with myc tag, pCMV-Myc | This study |
| HA-tUI, pET28 | Gift from Robert Cohen<br>Addgene plasmid #122662 |

## Acknowledgments

We thank B. Ebert (DFCI, Boston, MA) for the gift of HEK293T IKZF3 reporter cells and MM.1s Cas9 cells. We thank Z. Lou (Mayo, Rochester, MN) for the gift of RNF19A plasmids. We thank K. Shokat (UCSF, San Francisco, CA) for access to Xevos LC-MS and CytoFLEX. We thank Q. Zheng for help with LC-MS-MS. We thank A. Goldberg, D. Lee, B. Cravatt, W. Gibson, R. Puram, B. Ebert, M. Slabicki, T. Tian, D. Godage, and J. Gordan and Gordan lab members for helpful discussions and advice. Finally, we thank all members of the Ostrem, Schreiber, and Kadoch labs for helpful feedback and discussion throughout this project.

## Author contributions

Conceptualization: CK, SLS, JMLO. Investigation: WL, EGR, DCM, JMC, MM, JMK, GMM, VV, GMM, RS, JLP, MGR, JMLO. Funding acquisition: SPG, CSM, CK, SLS, JMLO. Supervision: CK, SLS, JMLO. Writing – original draft: WL, JMLO. Writing – review & editing: All listed authors

## Data and materials availability

All data are available in the main text or the supplementary materials. Correspondence and requests for materials should be addressed to JMLO

